# Dynamics of brain-muscle networks reveal effects of age and somatosensory function on gait

**DOI:** 10.1101/2023.02.02.526912

**Authors:** Luisa Roeder, Michael Breakspear, Graham K. Kerr, Tjeerd W. Boonstra

## Abstract

Walking is a complex motor activity that requires coordinated interactions between sensory and motor systems. We used mobile EEG and EMG to investigate the brain-muscle networks involved in gait control during overground walking in young, older and individuals with Parkinson’s Disease. Dynamic interactions between the sensorimotor cortices and eight leg muscles within a gait cycle were assessed using multivariate analysis. We identified three distinct brain-muscle networks during a gait cycle. These networks include a bilateral network, a left-lateralised network activated during the left swing phase, and a right-lateralised network active during right swing. The trajectories of these networks are contracted in older adults, indicating a reduction in neuromuscular connectivity with age. Individuals with impaired tactile sensitivity of the foot showed a selective enhancement of the bilateral network, possibly reflecting a compensation strategy to maintain gait stability. These findings provide a parsimonious description of interindividual differences in neuromuscular connectivity during gait.

**Teaser:** Dynamic network analysis shows how brain-muscle connectivity during gait varies with age and somatosensory function.

## Introduction

Human gait requires the activation and coordination of numerous muscle groups, adjusted on the fly to accommodate changing environmental and task demands. This rests upon the integration of sensory information, and the corresponding adaption of spinal and supraspinal motor commands (*1*). Spinal interneuron networks (central pattern generators) generate the basic locomotor pattern, which is modified by incoming peripheral afferent signals and descending supraspinal signals (*2*). Sensory afferent information contributes to corrective reflexes following sudden perturbations and may be used to adapt and update gait patterns (*3*). In humans the cortex not only contributes to complex walking tasks such as precision stepping or obstacle avoidance, but also to simple stereotyped locomotion. While the anatomy and physiology of the human sensorimotor system are well known, the functional organisation of the neural control of human gait remains incompletely understood.

Recent advances in mobile brain-body imaging (MOBI) now allow assessment of cortical processes during gait, in addition to muscle activity and 3D kinematics (*4, 5*). Using high-density electroencephalography (EEG), several studies have shown intra-stride modulations of oscillatory activity in the sensorimotor cortex, in particular in the beta band (*6-12*). Beta activity is a well-documented feature of the motor system during voluntary movement (*13*), and is likely involved in maintaining a particular sensorimotor state (*14-16*). During walking, beta oscillations have been found to be a sensitive marker for motor system engagement, with decreased beta power (event-related desynchronisation, ERD) just before and during the double support phase of the gait cycle and increased beta power during single-support and swing phases (*8, 9*).

Research on corticomuscular connectivity has shown that cortical beta activity is transmitted to spinal motoneurons during walking (*17-19*), particularly during the double support phase of the gait cycle (*20, 21*). This cortical input to spinal motor neurons appears to be reduced in the elderly and may be linked to changes in the temporal coordination of gait (*22*). Zandvoort and colleagues (*21, 23*) found that cortical beta-band activity was linked with specific patterns of distributed muscle activity (i.e. muscle synergies), which are involved in controlling shear forces to decelerate and accelerate the body (*24*). Indeed, it is thought that these cortical beta oscillations may be linked to postural stability during locomotion (*6*). For instance, mechanical perturbations to standing balance increased connectivity between the supplementary motor area and other central motor areas, while visual perturbations decreased connectivity between parietal and occipital regions (*25, 26*). These findings suggest that synchronized oscillatory activity has a functional role in how the central nervous system (CNS) coordinates behaviours such as gait (*27-29*), and may be particularly involved in the coordinating muscle activities for balance control.

A key challenge remains the high dimensionality of neurophysiological data across the gait cycle. While several studies have revealed interactions between specific muscles or areas of the CNS (*13, 14, 30*), the global organisation of the sensorimotor system remains elusive. Complex network analysis offers the opportunity to investigate the large-scale organisation of distributed systems (*31*) and the role of neural synchronisation in motor coordination (*32*). For example, network analysis has been used to map and analyse functional muscle networks during postural tasks, showing functional connectivity between different muscle groups at multiple frequency bands (*33, 34*) and between groups of spinal motor neurons (*35*). While complex network analysis allows mapping of interactions in large-scale systems, the dimensionality of these networks remains a challenge when comparing their topologies between experimental conditions or patient groups (*36, 37*). Moreover, functional networks are not static but reorganize and adapt to a task at different time scales (*38-40*). However, the dynamics in these large-scale networks generally unfold on a low-dimensional manifold or subspace (*41-44*). Exploiting these manifolds through appropriate dimension reduction techniques then renders the emergent network dynamics tractable for statistical comparisons.

Here we assess brain-muscle networks to investigate the distributed neural circuitry supporting locomotor behaviour and interindividual differences observed with healthy ageing and neuromuscular dysfunction. To achieve this, we estimate dynamic functional connectivity between bilateral sensorimotor cortices and eight leg muscles during overground walking in healthy young, older and people with Parkinson’s Disease (PD). Using unsupervised dimension reduction and graph theoretical analyses, we assess the dynamics of the brain-muscle network across the gait cycle within a low-dimensional subspace. The topology and time-frequency profiles of brain-muscle networks offer a unique window into the dynamics of the neural circuitry driving the musculoskeletal system during walking and how the connectivity dynamics varies with age and movement disorders.

## Results

### Time-frequency coherence

We estimated time-frequency coherence across gait cycles between eight leg muscles and left and right sensorimotor cortices. Intermuscular coherence (IMC) and corticomuscular coherence (CMC) revealed a distinct modulation within the gait cycle with peak values at frequencies <20 Hz during the double support phase (Figure 1). Strongest IMC was observed amongst muscles of the same leg (e.g., TA, SOL, GM, GL of the left leg), and particularly amongst triceps surae muscles. IMC was much lower between muscles of different legs. Overall, CMC was lower than IMC and mainly restricted to the double support phases of the gait cycle. Cortico-cortical coherence (CCC) was elevated throughout the gait cycle and showed minor increases during the double support phase. These time-frequency profiles were similar for all groups across all channel combinations (Figures S1 to S3), with coherence slightly reduced in older people and people with PD.

**Figure 1.**
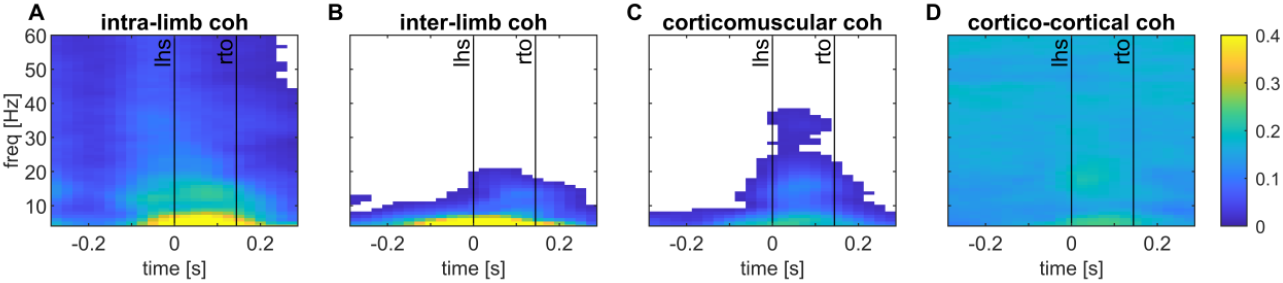
Time-frequency coherence between EMG and EEG channels during overground walking in healthy young people. Coherence is shown for a pair of muscles within the same leg, i.e., between the left Soleus muscle and the left lateral Gastrocnemius (**A**), between two muscles of each leg, i.e. between the left Soleus and right lateral Gastrocnemius (**B**), between the motor cortex and a contralateral muscle, i.e. between the left Soleus muscle and the right motor cortex (**C**), and between the left and right motor cortices (**D**). Coherence values are thresholded: average coherence values below the 95% CI are set to zero (white). The x-axis shows the time in seconds relative to heel strike (t=0) of the left foot and the y-axis the frequencies in Hz. Black vertical lines indicate the footswitch events (lhs, left heel strike; rto, right toe-off).

### Brain-muscle networks

To extract the low-dimensional subspace in which the brain-muscle networks unfold, we decomposed time-frequency coherence in canonical frequency bands using orthogonal NNMF. Three orthogonal components (networks) were extracted that explained most of the variance (72.7%). We used the scree plot criterion to determine this meaningful number of dimensions in our latent dimension-reduction analyses (Figure S4). Each component consists of the weights of a multilayer network and the temporal activation of that network within the gait cycle (Figure 2).

**Figure 2.**
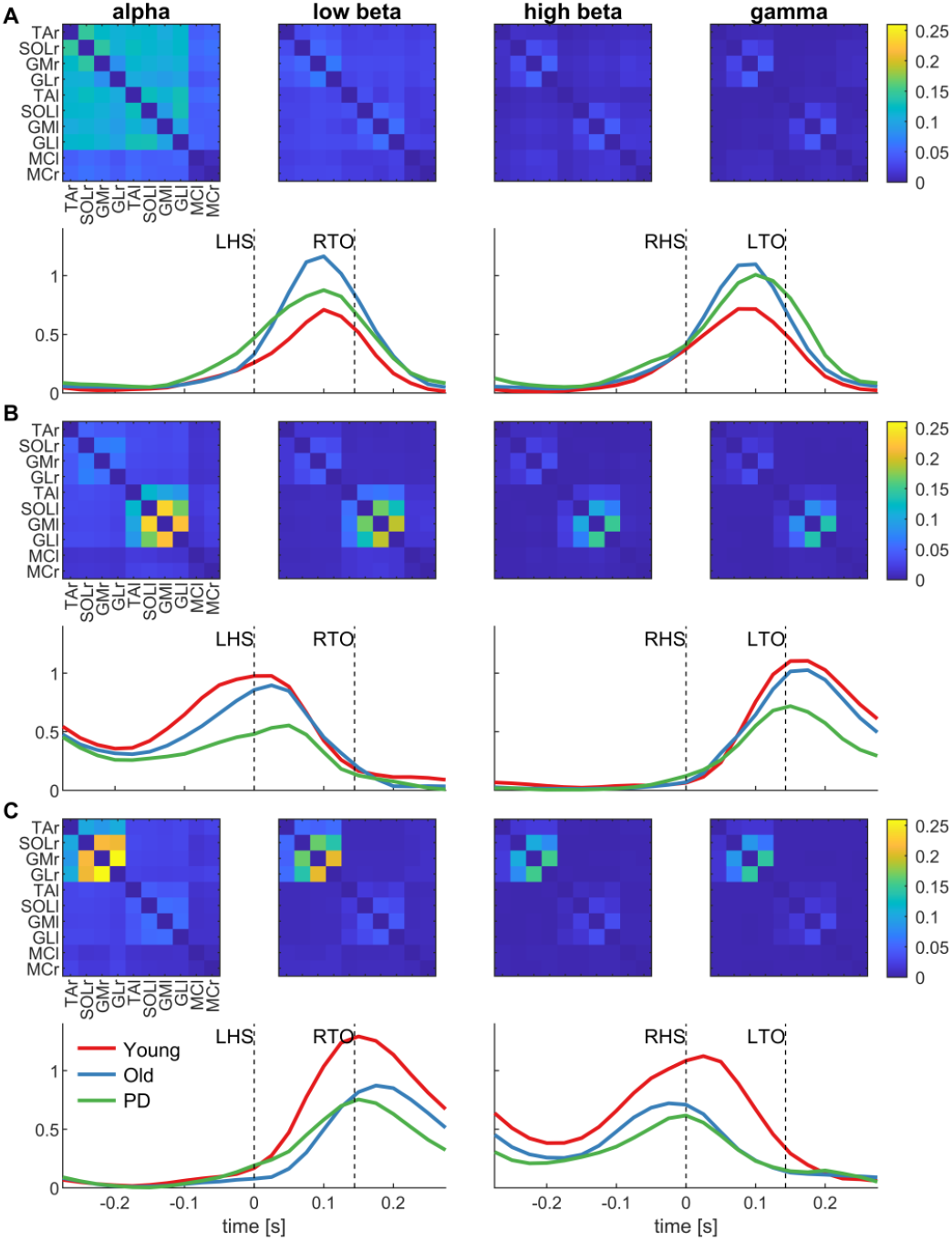
Network weights and activations obtained using orthogonal non-negative matrix factorisation of inter-/corticomuscular coherence. Coherence spectra of all EEG and EMG channel combinations, participants, conditions (left, right) and time points of the gait cycle are decomposed into three components (networks). Each network (**A, B, C**) is characterised by the weights (adjacency matrices) and activations time course throughout the gait cycle. The adjacency matrices show the average weights across participants for each frequency band separately (alpha, low beta, high beta, gamma). These weights give the edges between the 10 nodes of each network. The temporal activation patterns of each component are disaggregated by group (young, old, PD). LHS, left heel strike; LTO, left toe-off; MCl/MCr, motor cortex left/right; GMl/GMr, gastrocnemius medialis left/right; GLl/GLr, gastrocnemius lateralis left/right; RHS, right heel strike; RTO, right toe-off; SOLl/SOLr, soleus left/right; TAl/TAr, tibialis anterior left/right.

The first network (Figure 2A) shows strong connectivity between lower leg muscles within and across legs particularly at alpha frequencies, i.e., symmetric activation in the left and right leg. Temporal activation profiles reveal the strongest activation at middle of the double support phase when both feet are on the ground, which is symmetrical with respect to left and right heel strike. This network is only weakly activated during the swing phase. Visual inspection of the activation plots suggests higher activation in the older and PD groups; however, this effect is not statistically significant (F_2, 65_ = 1.23, p_adj_ = 0.30).

The second network (Figure 2B) also shows strongest connectivity at alpha and low beta frequencies with the strongest edge weights between muscles of the left leg. The temporal activation peaks when the left foot is raised, remain elevated during the swing phase of the left leg, and then peak again when the left foot hits the ground. Activation amplitude is lowest in the PD group compared to the young and old groups, but the group effect is not significant (F_2, 65_ = 2.81, p_adj_ = 0.10).

The third component (Figure 2C) is a right lateralised network with connectivity primarily in alpha and low beta frequencies. The temporal activation and network weights are a mirror image of network 2: the strongest edges are between muscles of the right leg and the network is activated from right toe-off to right heel strike, i.e., during the right leg swing phase. A one-way ANOVA revealed a significant group effect (F_2, 65_ = 6.88, p_adj_ = 0.006). Using post-hoc tests, we found that the young group had a significantly higher activation amplitude than the older (t(44) = 2.78, p_adj_ = 0.012) and PD group (t(40) = 3.43, p_adj_ = 0.004). There were no differences between the older and PD groups (t(42) = 0.56, p_adj_ = 0.58).

### Network topologies

To visualise the topologies of the three networks that we extracted, we thresholded the weights to obtain minimally-connected binary networks that include the different frequency bands as layers (Figure 3). The threshold was 0.145 for network 1, 0.278 for network 2, and 0.30 for network 3. Figure 3 shows how the thresholded, binarized networks are spatially distributed across the body (plotted separately per network and layer). Multilayer network analysis (*45*) was used to describe the identified networks; for these analyses we used the non-thresholded weights. Specifically, we calculated multilayer strength centrality for each network (bar graphs in Figure 3). The distribution of strength centrality shows that network 1 has a bilateral distribution with high weights across all muscles of both legs, which indicates that all muscles have similarly strong connectivity and are therefore equally important in the network. Network 2 shows peak weights at the triceps surae muscles of the left leg suggesting that these muscles have the strongest links within this network. Similarly, network 3 shows peak weights at the triceps surae muscles of the right leg.

**Figure 3.**
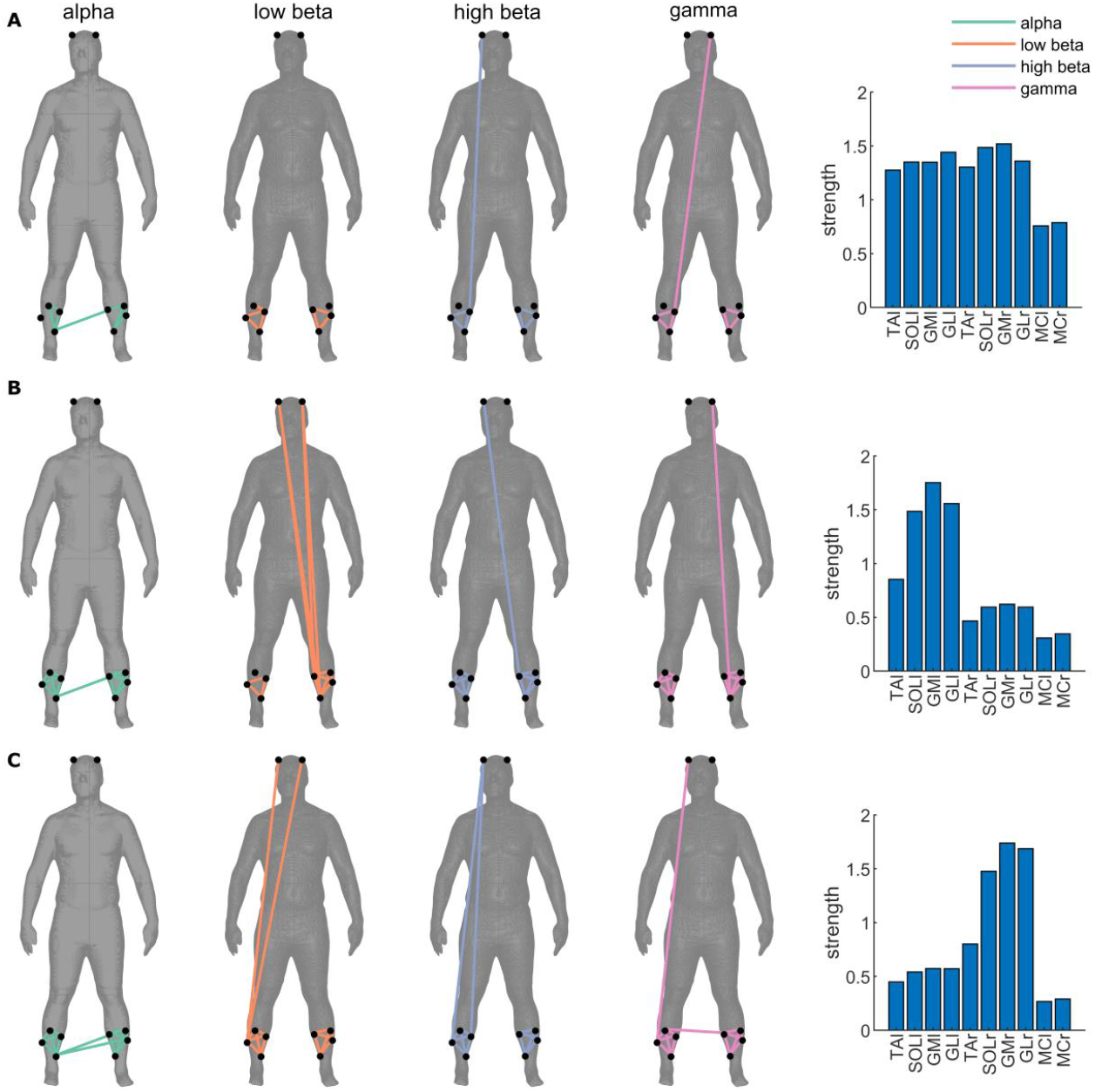
Topologies of binarized multilayer networks. Topologies displayed over human body show the minimally connected networks (thresholded edge weights; threshold was 0.145 for network 1 (**A**), 0.278 for network 2 (**B**), and 0.30 for network 3 (**C**)). The layers of each network (alpha, low beta, high beta) are displayed as separate columns and with different colour coding (connections at alpha (green), low beta (orange), high beta (blue) and gamma (pink) frequencies). The right-hand side panels show multilayer strength centrality for each network estimated using non-thresholded weights. Each bar graph shows the strength for each node (muscles or cortical sites) of each multilayer network. The higher the strength of a node, the stronger the connectivity with other nodes of the network. MCl/MCr, motor cortex left/right; GMl/GMr, gastrocnemius medialis left/right; GLl/GLr, gastrocnemius lateralis left/right; SOLl/SOLr, soleus left/right; TAl/TAr, tibialis anterior left/right.

### Low-dimensional subspace of network activations

The orthogonal nature of NNMF decomposition allows us to project the high-dimensional dynamic connectivity data onto the low-dimensional subspace spanned by the three networks. This reveals how the complex brain-muscle network follows a closed trajectory with the gait cycle that captures a continuous transition in relative dominance of each of these network modes (Figure 4A). That is, there is no serial activation of these networks but rather they are co-expressed, although each stage of the gait is associated with relative dominance of each of the networks.

**Figure 4.**
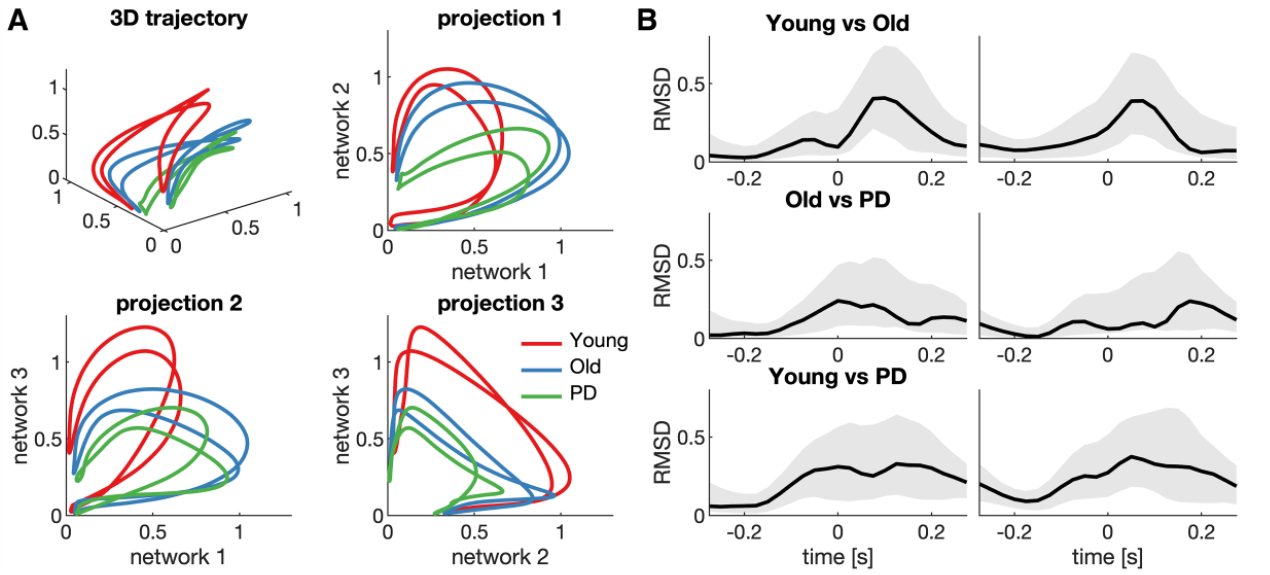
State-space embedding of the trajectories of the temporal activations of the three extracted networks. **A)** Different lines depict the group-average trajectories (Young, Old, PD). In addition to the trajectories in the 3D subspace, the projections on each of the three axis is shown. **B)** The root-mean-square deviation (RMSD) of the group-average trajectories of young vs old, old vs PD, and young vs PD. Grey-patches show the 95% confidence interval estimated based on 1000 bootstrapped samples.

The temporal order of activation of the three networks (i.e., network 1 during the middle of the double support phase, network 2 at left heel strike and network 3 at right toe off; see Figure 2) results in a rotation through in the three-dimensional subspace. The trajectories do not pass through the origin (0,0,0) as networks 2 and 3 remain activated during the left and right swing phase, respectively. The 2D projections show that while the younger group has a relatively stronger activation of the lateralized networks 2 and 3, the older and PD groups show relatively stronger activation of the symmetric network 1. The root mean square differences (RMSD) between the group-average trajectories show the phases of the gait cycle during which the trajectories differ most (Figure 4B). We used permutation testing to statistical test for group differences in the trajectories: individual participants were randomly swapped between the groups and the overall RMSD in the empirical trajectories were compared against these resampled null distributions. This shows that the trajectory of the young group differed from the old group (RMSD = 0.19, p = 0.046) and the PD group (RMSD = 0.24, p = 0.017), while there was no significant difference between trajectories of old and PD groups (RMSD = 0.12, p = 0.24).

### Inter-subject variability in network activations

We then performed principal component analysis (PCA) to further assess the inter-subject variability in the network activations in the 3D subspace (*46-48*). The first principal component (PC1) explained 41% of the inter-subject variability in the 3D trajectories and reflected the average pattern of the 3D trajectories: bilateral network 1 peaked in the middle of the double support phase, left-lateralized network 2 peaked at left heel strike and toe off, and right-lateralized network 3 peaked at right heel strike and toe off (Figure 5, top row). The eigenvector coefficients showed a significant group effect (F_2,63_ = 4.27, p = 0.017). Post-hoc test showed that the younger group had higher coefficients than the PD group (t(40) = 2.89, p_adj_ = 0.019), but no difference between the younger and older groups (t(44) = 1.33, p_adj_ = 0.19) or between the older and PD groups (t(42) = 1.86, p_adj_ = 0.10). As this component reveals large excursions in all three dimensions, the group effect captures a general reduction of network activation, that is, a shrinking of the 3D trajectories. The second component (PC2) explained 24% of the inter-subject variability and revealed a selective activation of the bilateral network 1 (Figure 5, bottom row). Although the eigenvectors coefficients were numerically higher in the old and PD groups, the group effect was not statistically significant (F_2,63_ = 2.18, p = 0.12).

**Figure 5.**
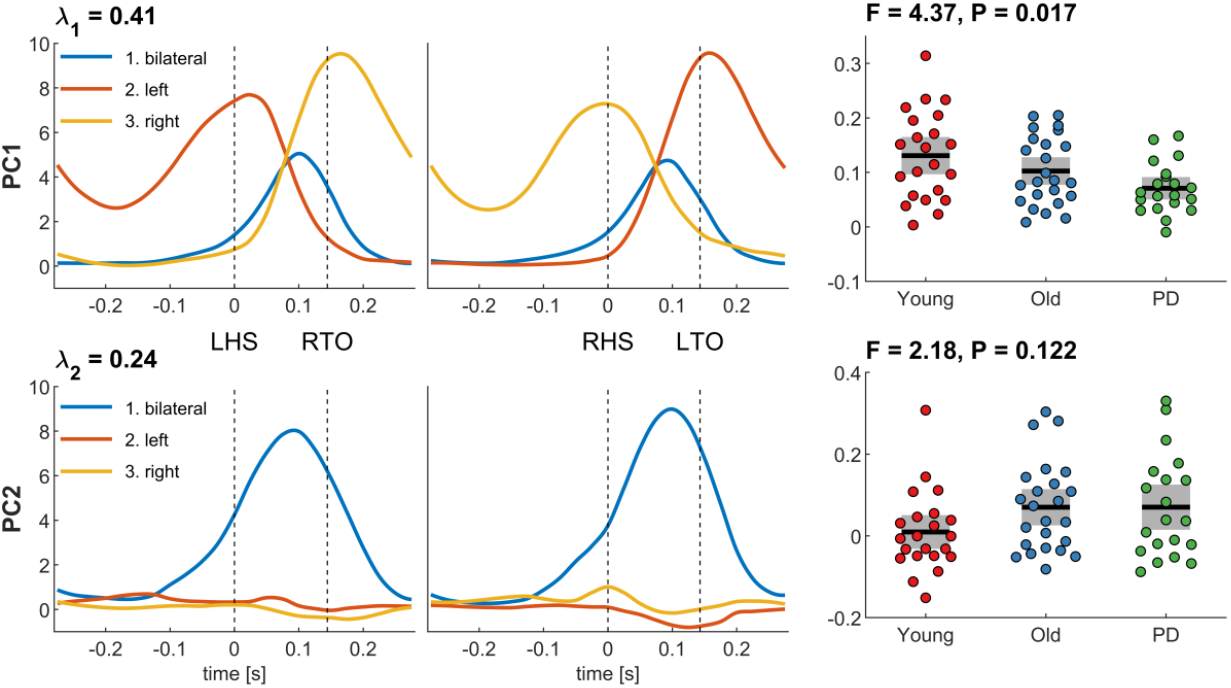
Principal component analysis and group effects of network activations. For both principal components (PC1 and PC2) we show the network activations throughout the gait cycle (left column) and the eigenvector coefficients per group (right column). Coloured dots show individual data of each participant, black horizontal lines show the group mean, grey boxes show the SEM. PC, principal component; λ, explained variance; LHS, left heel strike; RTO, right toe-off; RHS, right heel strike; LTO, left toe-off.

Finally, we explored the association between the eigenvector coefficients and several participant characteristics (age, sex, weight) and clinical scores (touch sensitivity, UPDRS motor sub-score, Hoehn&Yahr). When controlling the false discovery rate, we found a significant correlation between PC1 and age (r = 0.33, p_adj_ = 0.042, n = 66) and between PC2 and touch sensitivity (r = -0.41, p_adj_ = 0.042, n = 43; see Figure S5 and Table S1). The correlation effect between PC1 and age is negative showing that PC1 decreases at older age. The correlation between PC2 and touch sensitivity is also negative indicating that in older people with and without PD (touch sensitivity data only available in old and PD groups) PC2 is reduced as touch sensitivity deteriorates.

## Discussion

We identified three distinct brain-muscle networks underlying human gait: 1) a bilateral network that is left-right symmetric and mostly active during the middle of the double support phase, 2) a left-lateralised network that is activated during the left swing phase and peaks at the left heel strike and toe-off, 3) a right-lateralised network that mirrors the left-lateralised network. The three networks span a low-dimensional subspace in which neuromuscular gait dynamics unfold. These trajectories reveal that these networks are simultaneously co-expressed during the double support phase but with different networks dominating the initial and terminal double support phase. The spread of these trajectories through their low-dimensional subspace exhibits an age-related contraction, corresponding to a general reduction in neuromuscular connectivity at older age and a selective increase in activation of the bilateral network in participants with impaired tactile sensitivity of the foot sole.

The current findings provide a parsimonious description of neuromuscular connectivity during gait in three spatiotemporal patterns. These three patterns have similar spectral profiles – with strongest coherence in the alpha and lower beta band – and are all primarily recruited during the double support phase. However, these modes have distinct network topologies and temporal profiles that are aligned to specific behavioural events. These findings demonstrate that neuromuscular connectivity followed a low-dimensional manifold. By assessing dynamic brain-muscle networks, we show that neuromuscular connectivity is not shaped by a simple cortical drive to the muscle but reveals distinct network motifs that likely reflect different neural circuits. These circuits – rather than alternating – perform a low-dimensional rotation in neural space, in which the connectivity is continuously cycling through different network configurations within the gait cycle, which bears resemblance to the ensemble dynamics observed in spinal motor networks (*49*). This low-dimensional rotation may reflect a spiral attractor of the spinal interneuron network, in which changes in the processing of afferent or efferent activity may shape the underlying manifold and therefore change the attractor (*50, 51*).

Previous research has shown increased beta activity during the double-support phase in cortical activity (*6-9*) and corticomuscular coherence (*20, 21, 23*), suggesting a corticospinal drive to the muscles (*18, 52*). Here we show the modular organization of common spinal input to different lower leg muscles, which compliments previous notions of modular control of gait (*24, 53, 54*). In particular, muscle synergy analysis has revealed two lateralized muscle synergies that are involved in the touch-down and lift-off of the left and right legs, which show coherent activity with the motor cortices (*21, 23*) and emerge around the onset of independent walking in children (*24*). The two lateralized brain-muscle networks identified in the current study were also precisely timed to the critical events of the gait cycle – toe-off and heel strike – and remained activated while the foot was in the air. It appears therefore likely that these networks are involved in stabilising the ankle joint rather than in generating forward propulsion, in line with the suggested role of beta activity maintaining sensorimotor states (*27*).

PCA of the network activations showed that the first principal component captures a general decline (contraction/shrinking) in activation of all three networks between groups (Young > Old > PD) and is negatively correlated with age. That is, the first component was reduced in older people with and without PD. Overall, this reflects a general reduction in neuromuscular connectivity with older age. This finding agrees with previous studies showing decreased corticomuscular and intramuscular coherence during walking in older people (*19, 22, 55, 56*) or in people with PD (*57*), which may be indicative of decreased sensory processing in older people (*58*). Older people exhibit reduced reactive locomotor adjustments to perturbations (*59, 60*) and particularly temporal aspects of gait control are affected by age, e.g., double support time (*61*). Such reactive gait adaptations are mainly driven by proprioceptive and visual feedback, which is processed by spinal circuits (*58*), and involved in modifying reactive changes in muscle activity (*62*). Importantly, spinal reflex pathways appear to be affected by age, resulting in reduced amplitude of the H-reflex (*63*) and reduced modulation of the H-reflex (*64*) during walking. Together this suggests that the general reduction in neuromuscular connectivity may reflect reduced input from spinal circuits into lower leg muscles that are involved in temporal gait control.

In addition to the general reduction in neuromuscular connectivity, the second principal component (PC2) revealed a selective modulation of the bilateral network 1, which was enhanced in the older and PD groups. We found a significant negative correlation between PC2 and touch sensitivity scores. That is, PC2 is higher in older and PD people, and within this group of older people the component is reduced as touch sensitivity deteriorates. This indicates that older people with and without PD use more of the bilateral network to control their gait rather than the left and right lateralised networks. This strategy may reflect a preference for a more stable, symmetric gait pattern in older people, particularly when touch sensitivity is impaired. Kinematic studies found older people show a reduced medio-lateral shift of the centre of mass (COM) – a strategy which is more robust towards perturbations and prioritizes maintaining mediolateral balance – compared to young adults who shift their body centre away from the fast foot during split-belt treadmill walking, leading to an asymmetric balance control that may be metabolically more efficient (*65, 66*). Indeed, older people who are destabilised through visual perturbations increase step width to gain stability (*67*). The selective activation of the bilateral network in older participants may be linked to a reduction in muscle spindle afferent information relevant for the control of medio-lateral gait stability (*68*) or a decrease in sensitivity of cutaneous mechanoreceptors of the feet (*69, 70*). Receptors in the skin around the ankle and foot sole may provide information about ankle rotation (*71*) and influence foot trajectory variability during walking (*72*). Decreased sensory information from skin receptors and muscle spindles in older people likely affects foot placement and stepping strategy, and thereby affects mediolateral balance control. Hence, the observed increase of the bilateral network 1 may reflect a compensation strategy to maintain mediolateral gait stability in response to impaired sensory processing at the feet.

As the primary aim of our study was descriptive, the explanatory or causal interpretations of the possible neuronal origins of these three spatiotemporal patterns should be made with caution. Recording 3D kinematics would allow assessment of the link between neural activity and locomotor behaviour more directly, for example by manipulating mediolateral stability through experimental perturbations (*66, 67*). In addition, the limited number of EEG channels available did not allow us to perform a source reconstruction of the signals, which would have provided a clearer neurophysiological understanding of the current results and delineate the cortical regions involved in the neural control of gait. While we estimated imaginary coherence between sensors over bilateral sensorimotor cortex (*73*), functional connectivity in channel space should be interpreted cautiously as the channel locations cannot be seen as an approximation of a source’s anatomical location and spurious connectivity may occur between sensors (*74*). Finally, EMGs were only recorded from lower leg muscles, and acquiring data from a higher number of EMG channels from leg, hip and arm muscles will provide a more complete picture of brain-muscle network modular gait control. For example, directed connectivity analyses revealed that upper limb muscles drive and shape lower limb muscle activity during gait via subcortical and cortical pathways (*75*). Future work could also apply the current network framework to gait perturbation paradigms that require locomotor adjustments, to test if the bilateral network indeed reflects a compensation strategy to maintain mediolateral gait stability.

## Materials and Methods

Three groups participated in this study with 69 participants in total: 1) 24 healthy young adults (12 females, mean (SD) age 25.9 (3.2) years), 2) 24 healthy older adults (12 females, mean (SD) age 65.1 (7.8) years), 3) 21 individuals with PD (8 females, mean (SD) age 67.4 (7.3) years). Participants were screened for visual and musculoskeletal injuries/diseases, neurotropic medication and injuries/disease of the central nervous system (e.g., spinal cord injury, MS, neuropathies) in order to determine eligibility. All experimental protocols were approved by the Human Research Ethics Committee of Queensland University of Technology (#1300000579) in accordance with the Declaration of Helsinki, and all participants gave written informed consent prior to participation. These data were previously reported in Roeder et al. (*22*).

### Experimental protocol

Participants performed overground walking for 12-14 minutes. Participants walked barefoot at their preferred speed (3.3 – 4.8 km h^-1^) with natural arm swing (mean (SD) gait velocity in km h^-1^ per group: Young 4.17 (0.41), Older 4.06 (0.49), PD 4.06 (0.50)). Participants walked back and forth along a straight path (∼14 m) on a firm surface in the gait laboratory and turned at each end of the room. The turning sections, including acceleration at the beginning of the straight-line path and deceleration at the end (∼2.4 m), were excluded from further analyses and only straight-line walking (8.9 m) was used for analysis.

Healthy older people and people with PD also completed a peripheral sensation test on a different day. We assessed tactile sensitivity at different sites of their feet (lateral malleolus, plantar surfaces of the great toe, midfoot/arch and heel) using Semmes-Weinstein pressure aesthesiometer with nylon monofilaments (*76*). People with PD were also assessed for disease severity using the Unified Parkinson’s Disease Rating Scale (MDS-UPDRS) and Hoehn & Yahr scoring (*77, 78*). People with PD presented in their optimally medicated state for all experiments (group mean (SD) levodopa equivalent daily-dose 544 (241) mg).

### Data acquisition

EEG, EMG and kinematic events (heel strike and toe off) were recorded while participants performed overground walking. Bipolar surface EMG was recorded from the left and right tibialis anterior (TA), soleus (SOL), medial and lateral gastrocnemii (GM, GL) muscles. Simultaneously, EEG signals were recorded using water-based AgCl electrodes placed at 10 cortical sites according to the international 10-20 standard (P3, P4, C3, Cz, C4, F7, F3, Fz, F4, F8). Footswitches were attached onto the participants’ sole at the heel and the big toe of both feet. All signals (EEG, EMG and footswitches) were recorded synchronously with a wireless 32-channel amplifier system (TMSi Mobita, The Netherlands) and sampled at 2 kHz. The recording system and wireless transmitter were placed in a belt bag and tied around the participants’ waist.

### Data pre-processing

EEG, EMG and footswitch data were pre-processed as reported in (*22*). In short, electrophysiological data were normalized to unit variance and segments with excessive noise (large-amplitude movement artefacts, EMG activity) were removed. Due to excessive EEG artefacts across all channels and conditions (large-amplitude movement artefacts (>300μV) occurring with regular rhythmicity in each gait cycle throughout the entire record) we excluded two healthy young participants and one participant with PD. Hence, analyses were based on n = 22 for the younger group, n = 24 for the older group, and n = 20 for PD group.

EEG signals were band-pass filtered (2^nd^ order Butterworth, 0.5-70 Hz) and re-referenced to a common average reference (EEGLAB version 13.6.5, *79*). Independent components containing eye blink, muscle, or movement artefacts were removed from the data. Finally, we calculated bipolar EEG montages to assess cortical activity from bilateral sensorimotor cortices: C3-F3 for the left sensorimotor cortex, and C4-F4 for the right sensorimotor cortex (*80*).

EMG data were high-pass filtered (4^th^ order Butterworth, 20 Hz cut-off) and full-wave rectified using the Hilbert transform (*81*). For coherence analysis EMG signals were also demodulated to avoid spurious coherence estimates resulting from periodic changes in EMG amplitude (*82*).

### Time-frequency analysis

We used time-frequency analysis to assess changes in spectral measures within the gait cycle (*83*). EEG and EMG signals were segmented into 220 segments of 600 ms length (−300 to +300 ms with respect to heel strike). Heel strike served as reference point (t = 0) in the gait cycle to which all segments were aligned. Event-related coherence spectra were estimated across the 220 segments using Short-time Fourier transform (a 375-ms Hanning window with increments of 25 ms) by squaring the absolute value of coherency. We computed coherence pairwise between all possible channel combinations (EMG-EMG or intermuscular coherence; EEG-EMG or corticomuscular coherence; EEG-EEG or corticocortical coherence). To minimize the effect of volume conduction, we used imaginary coherence to estimate corticocortical connectivity between bilateral sensorimotor cortices (*84*). With a total of 10 channels (8 EMG and 2 bipolar EEG), we hence assessed dynamic functional connectivity between 45 channel pairs. Event-related coherence spectra were computed for each participant during overground walking.

### Network analysis

For dimension reduction of these data, we used orthogonal non-negative matrix factorisation (NNMF, *85*) to decompose coherence spectra across all channel combinations, time points of the gait cycle and participants into gait-cycle dependent activation patterns and their corresponding weights (i.e. coupling strength). These weights define the undirected weighted network for each frequency layer that is common across participants. In contrast, the activation patterns of these networks across the gait cycle vary between participants. We used an open-source MATLAB toolbox for computing the orthogonal NNMF (*86*), which is similar to non-negative sparse PCA (*86, 87*).

Weights were thresholded and binarized to obtain minimally connected networks for visualisation purposes, as described in Kerkman, Daffertshofer, Gollo, Breakspear and Boonstra (*34*). We used the Brain Connectivity Toolbox implemented in MATLAB to threshold the corresponding matrices (*36*). This yields a single, unique value, which corresponds to the percolation threshold, resulting in sparse networks in which each node is connected to at least one other node by an edge at one of the layers of the multilayer network. That is, connectivity at each frequency band (alpha, low beta, high beta, gamma) represents a layer of the multilayer network. To describe the multilayer networks further, we calculated multilayer strength centrality (*88*) using the R package MuxViz (v3.1, *45, 89*). For this purpose, we treated the layers of each network as non-interconnected (i.e., a network has multiple layers, which encode a specific relationship between nodes, and nodes preserve their identity across layers but are not connected across layers) and we used non-thresholded weights. Multilayer strength centrality is the sum of weights of edges connected to a node added up across all layers (*45*). Multilayer strength centrality quantifies the importance of specific nodes (i.e., muscles and cortical regions in this study) within a network: higher weights at a specific node indicate stronger connectivity to other sites/regions (and hence, stronger interactions). By looking at the distribution of strength centrality across the nodes (i.e., muscles and cortical regions) it is possible to identify which nodes are ‘key players’ in a network and which ones only contribute weak links.

To further investigate the dynamics of brain-muscle connectivity and its temporal evolution throughout the gait cycle, we assessed the state space trajectories in the low-dimensional subspace (*90*). This was possible because the networks we extracted via NNMF were orthogonal. To visualise these results, we projected the mean state space trajectory for each group (young, old, PD) onto a 3D plot.

### Statistical analysis

To compare the activation patterns of these networks across groups, we extracted the network activation amplitude by averaging the activation pattern across all time points of the gait cycle for each participant and network separately. These activation amplitude values were contrasted across groups (young, old, PD) using three separate one-way ANOVAs (one for each network). We controlled the false discovery rate by using the Benjamini-Hochberg procedure (*91*) using an opensource Matlab script (*92*).

To investigate the inter-subject variability in the network activations in the 3D subspace, we performed principal component analysis (PCA; *46, 47, 48*). The 3D trajectory of each participant was reshaped into a vector and PCA was performed across all participants (n=66), which decomposed this data into principal components (PCs) and the eigenvector coefficients that capture the contribution of each participant. We then compared the eigenvector coefficients of PC1 and PC2 between groups using a one-way ANOVA and explored the association with participant characteristics (age, sex, weight) and clinical scores (touch sensitivity, UPDRS motor sub-score, Hoehn&Yahr) using correlation analysis. We controlled the false discovery rate for these 2 × 6 = 12 correlations using the Benjamini-Hochberg procedure (*91*).

The significance level (alpha) was set at 0.05 for all statistical analyses. We used student t-tests as post-hoc analysis when the group effect was significant.

## Acknowledgements

Tjeerd Boonstra was supported by the European Union’s Horizon 2020 research and innovation programme under the Marie Sklodowska-Curie grant agreement No 895914. The authors declare that they have no competing interests.

All data needed to evaluate the conclusions in the paper are present in the paper and/or the Supplementary Materials.

## Author contributions

Conceptualization: LR, TWB, MB, GK. Data curation: LR. Formal analysis: LR, TWB. Investigation/Data collection: LR. Methodology: LR, TWB, MB, GK. Project administration: LR. Resources: GK. Software: TWB, LR. Supervision: TWB, GK. Visualization: TWB, LR. Roles/Writing - original draft: LR. Writing - review & editing: LR, TWB, MB, GK.

## Notes

### Competing Interest Statement

The authors have declared no competing interest.

### Summary of Updates

All parts of the manuscript were revised.

