## Supplementary Material for "Dynamics of brain-muscle networks reveal effects of age and somatosensory function on gait"

Luisa Roeder* *et al.*

**This PDF file includes:**

Figs. S1 to S5
Table S1

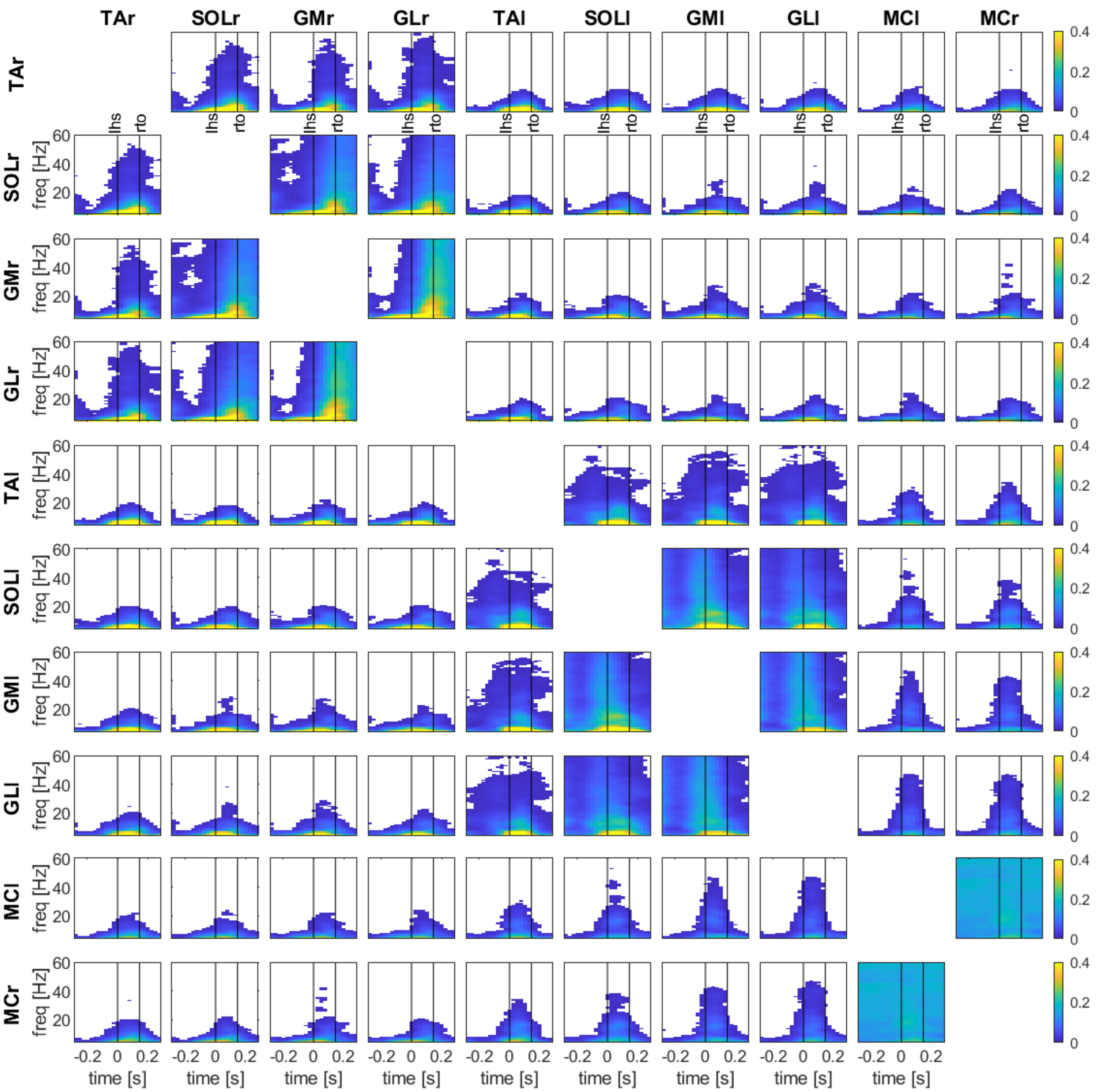


**Fig. S1.**

**Time-frequency coherence between all channel combinations during overground walking in healthy young people**. Coherence is shown between all four leg muscles (TA, SOL, GM, GL) on both sides (left indicated with l, right with r) and motor cortical sites (MCl, MCr). Coherence values are thresholded: average coherence values below the 95% CI are set to zero (white). The x-axis shows the time in seconds relative to heel strike (t=0) of the left foot and the y-axis the frequencies in Hz. Black vertical lines indicate the footswitch events. lhs, left heel strike; rto, right toe-off.

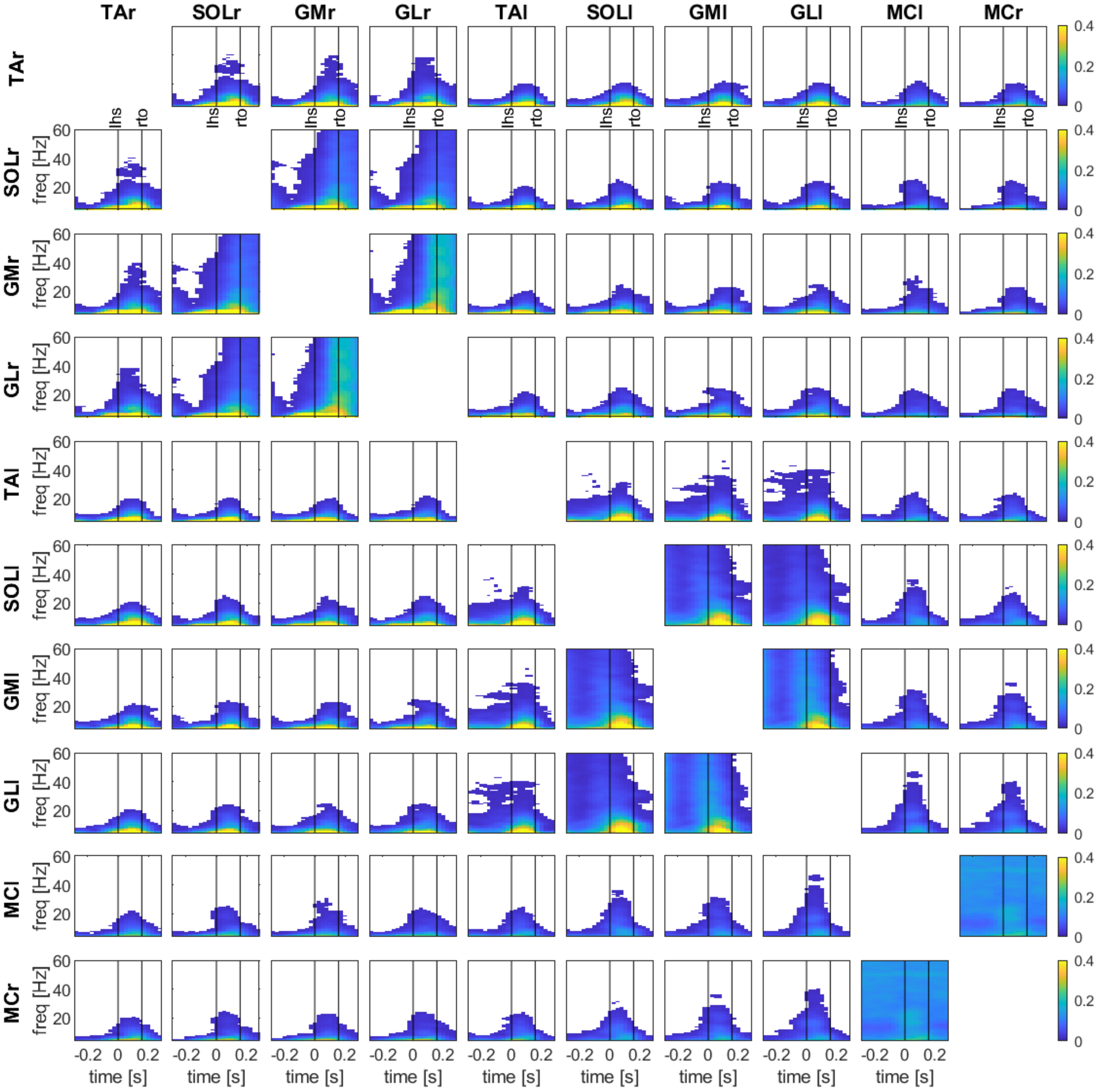


**Fig. S2.**

**Time-frequency coherence between all channel combinations during overground walking in healthy older people**. Coherence is shown between all four leg muscles (TA, SOL, GM, GL) on both sides (left indicated with l, right with r) and motor cortical sites (MCl, MCr). Coherence values are thresholded: average coherence values below the 95% CI are set to zero (white). The x-axis shows the time in seconds relative to heel strike (t=0) of the left foot and the y-axis the frequencies in Hz. Black vertical lines indicate the footswitch events. lhs, left heel strike; rto, right toe-off.

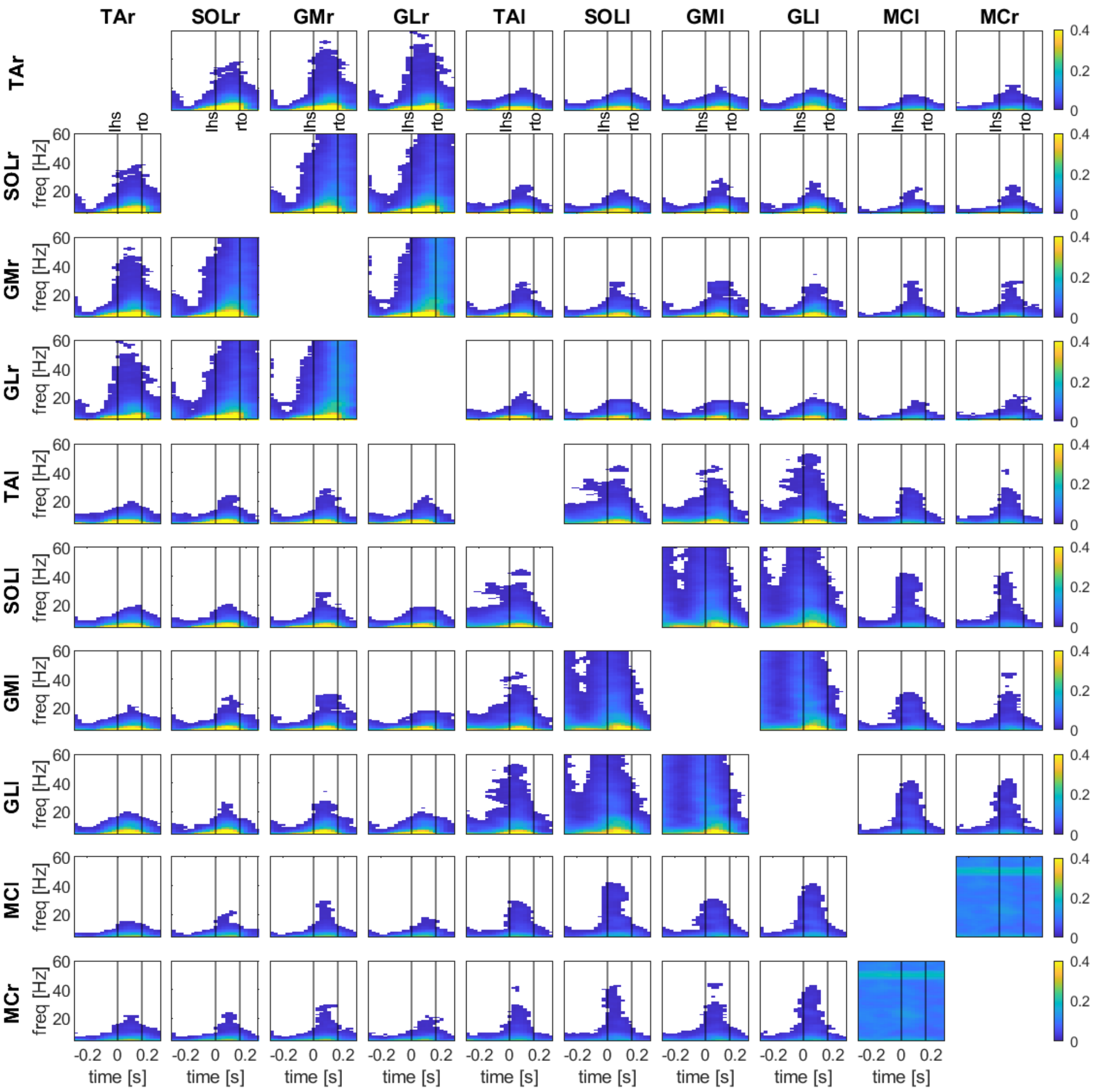


**Fig. S3.**

**Time-frequency coherence between all channel combinations during overground walking in people with PD**. Coherence is shown between all four leg muscles (TA, SOL, GM, GL) on both sides (left indicated with l, right with r) and motor cortical sites (MCl, MCr). Coherence values are thresholded: average coherence values below the 95% CI are set to zero (white). The x-axis shows the time in seconds relative to heel strike (t=0) of the left foot and the y-axis the frequencies in Hz. Black vertical lines indicate the footswitch events. lhs, left heel strike; rto, right toe-off.

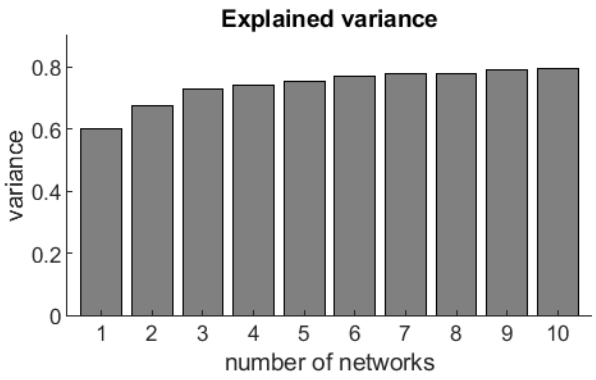


**Fig. S4.**

**Variance explained by components (networks).** Cumulative of the variance explained by 10 networks. Networks were extracted by orthogonal non-negative matrix factorisation (NNMF). Three networks explained 72.7% of the variance.

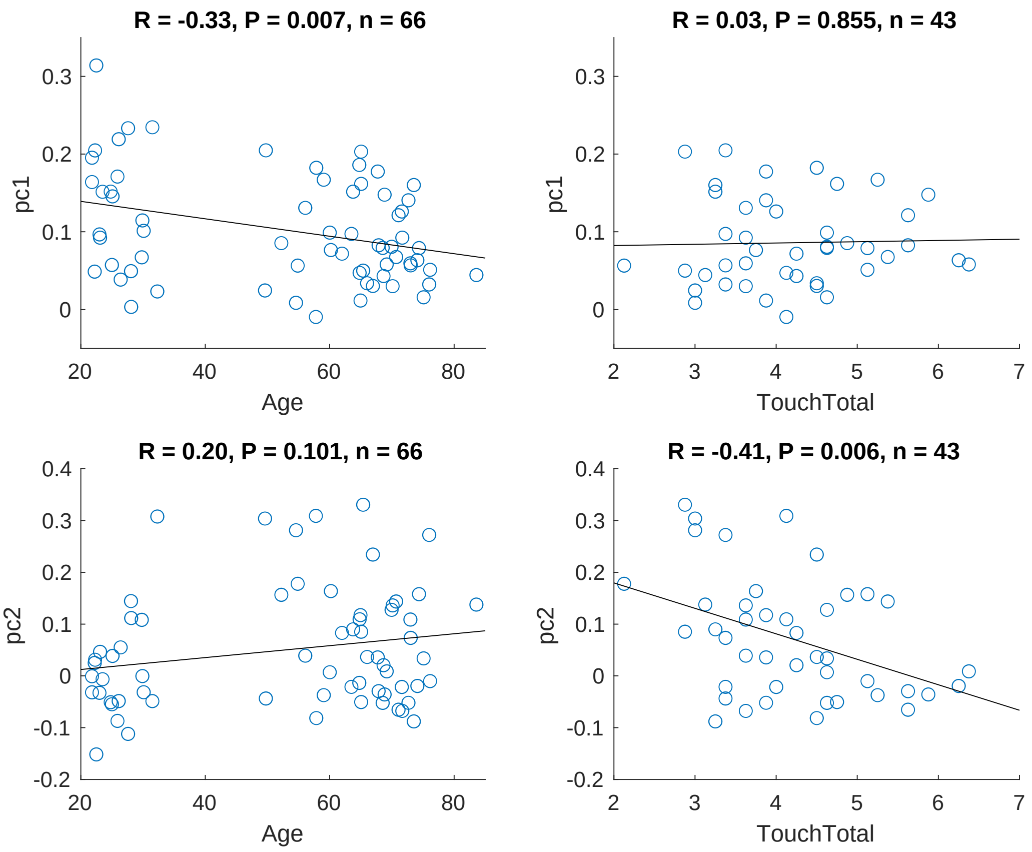


**Fig. S5.**

**Correlation effects of network activations and participant & functional measures.** Higher "TouchTotal" scores (touch sensitivity) indicate worse touch sensitivity.

| | PC1 | | | PC2 | | |
|---|---|---|---|---|---|---|
| | r | p | $p_{ajd}$ | r | p | $p_{adj}$ |
| Age | **-0.33** | **0.007** | **0.042** | 0.20 | 0.10 | 0.30 |
| Sex | 0.17 | 0.16 | 0.39 | -0.28 | 0.025 | 0.10 |
| Weight | -0.11 | 0.38 | 0.56 | -0.12 | 0.33 | 0.56 |
| $TouchSensitivity_{total}$ | 0.03 | 0.86 | 0.86 | **-0.41** | **0.006** | **0.042** |
| $UPDRS_{motor}$ | -0.23 | 0.33 | 0.56 | -0.09 | 0.72 | 0.85 |
| Hoehn&Yahr | -0.17 | 0.47 | 0.62 | 0.07 | 0.78 | 0.85 |

**Table S1.**
Correlation between the eigenvector coefficients and participant characteristics and clinical scores.
